# Control of AMPA receptor activity by the extracellular loops of auxiliary proteins

**DOI:** 10.1101/140053

**Authors:** Irene Riva, Clarissa Eibl, Rudolf Volkmer, Anna L. Carbone, Andrew J. R. Plested

## Abstract

At synapses throughout the mammalian brain, AMPA receptors form complexes with auxiliary proteins, including TARPs. However, how TARPs modulate AMPA receptor gating remains poorly understood. We built structural models of TARP-AMPA receptor complexes for TARPs γ2 and γ8, combining recent structural studies and de novo structure predictions. These models, combined with peptide binding assays, provide evidence for multiple interactions between GluA2 and variable extracellular loops of TARPs. Substitutions and deletions of these loops had surprisingly rich effects on the kinetics of glutamate-activated currents, without any effect on assembly. Critically, by altering the two interacting loops of γ2 and γ8, we could entirely remove all allosteric modulation of GluA2, without affecting formation of AMPA receptor-TARP complexes. Likewise, substitutions in the linker domains of GluA2 completely removed any effect of Y2 on receptor kinetics, indicating a dominant role for this previously overlooked site proximal to the AMPA receptor channel gate.

## Introduction

Since the identification of the protein Stargazin, also known as γ2, as the prototype transmembrane AMPA receptor regulatory protein (TARP)(1), a broad family of auxiliary proteins for the AMPA receptor have been described (2, 3). These proteins play an essential role in tethering AMPA-type glutamate receptors at the synapse, and also exert complex control over surface expression of functional receptors (4, 5). Auxiliary proteins regulate the function of AMPA receptors, with both positive and negative modulation of gating (6–9), as well as control over permeation and block (10). The range of auxiliary subunit influence over synaptic transmission is compounded by striking regional and cell-type specific expression (11, 12), and a patchwork of interaction patterns (13).

TARPs and other auxiliary proteins modify the gating and pharmacology of synaptic AMPA receptors (14, 15). The physiological importance of modulation is likely to be the specialization of particular codes of short-term plasticity, in the hippocampus and cerebellum at least (7, 16–18). Recently, antagonists of AMPA receptors that target GluA2–γ8 complexes were described (19, 20), further enhancing interest in the molecular basis of complexes of GluA subunits and their auxiliary proteins as potential drug targets.

Previous studies showed that some of the effects of auxiliary proteins on receptor gating were due to the extracellular domains (21–23). However, several of these studies made use of chimeras with γ5, which was presumed to be a null subunit, but which was subsequently shown to modulate gating and conductance of GluA receptors (24). Although some mutations in extracellular portions of TARPs were reported that affect TARP activity, there is no clear indication that these TARPs formed complexes with GluA subunits as well (25). On the other hand, some studies of assembly made use of functional tests to assess the strength of interaction (26). Given the variable stoichiometry of assembly between different TARP isoforms (27, 28), interpreting these data, which combine the strength of association, expression and modulation into a single metric, is difficult. Very recently, a chimeric approach confirmed impressions from structural studies that transmembrane interactions are important for proper assembly, with the TM3 and TM4 segments of γ2 and the M1-M3 helices of the AMPA receptor determining complex assembly. However, the C-termini of both the AMPA receptor and TARPs also appear to be involved (29). Despite these insights, there is very little information about the extent to which different domains contribute to gating of complexes (30), and no information about the structural basis of slow modulation, superactivation (37).

Two of the predominant TARPs in the brain are the auxiliary proteins γ2 and γ8. In this work, we isolate the extracellular segments of γ2 and γ8 that are responsible for modulation of gating, and show that these segments act on the receptor via the linkers connecting the ligand binding domain (LBD) and the transmembrane domain (TMD). In so doing, we were able to produce “null” TARPs, which assemble normally but show no modulation of gating. Hereby, we establish mechanisms for the subunit specific modulation of AMPA receptors by auxiliary proteins.

## Results

### A model of auxiliary protein interactions

Previous studies of TARP modulation of AMPA receptors have identified extracellular regions as potential interaction motifs. Crystal structures of Claudins, proteins with close homology to TARPs, enabled a more refined view, defining a folded extracellular “cap” (31–33) that substantially limits the sections of the extracellular portion of TARPs that are able to interact with the AMPA receptor, and therefore the likely range of these interactions. More recently, CryoEM micrographs of GluA2-TARP complexes allowed unambiguous positioning of TARPs at the periphery of the GluA2 pore, and partially resolved the extracellular domains of TARPs (34, 35). The major sequence and structural differences between Claudin and TARP proteins, and between TARPs with different modulatory effects, are found in the variable extracellular loops between β1 and β2 (Loop 1), and between TM3 and β5 (Loop 2). We sought to identify interactions between TARPs and the extracellular regions of the GluA2 receptor on this basis.

To understand the scope of TARP interactions with the AMPA receptor, we began by modeling the loops of γ2 and γ8 into a hybrid structure composed of Claudins and GluA2. Comparing these hybrid complexes to CryoEM electron density maps suggested that a range of interaction sites with the LBD-TMD linkers and D2 domains of the LBD are possible (Figures 1A and Figure 1 – Figure supplement 1A). Whereas TARP loop 2 (L2) engages in the receptors pore four-fold symmetry, loop 1 (L1) reaches up to the two-fold symmetry of the LBD layer. In other words, while L2 can interact four times in the same way with the receptor (Figure 1 – Figure supplement 1B), L1 has at least two distinct modes of interaction depending on to which receptor subunits the TARP is adjacent (subunit A-D and B-C, Figure 1A, or A-B and C-D, Figure 1 – Figure supplement 1A). The variable loop 1 is not resolved in structures to date, consistent with it being a flexible modulatory element. Superactivation of GluA2 receptors resembles strongly the slow modulation of AMPA receptors by particular allosteric modulators that bind at the dimer interface (36, 37). We reasoned that extracellular loop interactions that stabilized the superactive state could preferentially target the GluA2 LBD dimer. To test this hypothesis, we composed an overlapping library of hexameric peptides based on extracellular sections of TARPs, targeting the long loop 1 of γ2 and γ8, and other potential interacting sites (Figure 1 – Figure supplement 1C). Because the active dimer of LBDs ought to be intact for superactivation, we compared the interactions of our peptide library between the GluA2 LBD (flip form) and LBDs harboring the L483Y substitution, which greatly increases dimer formation in solution.

**Figure 1.**
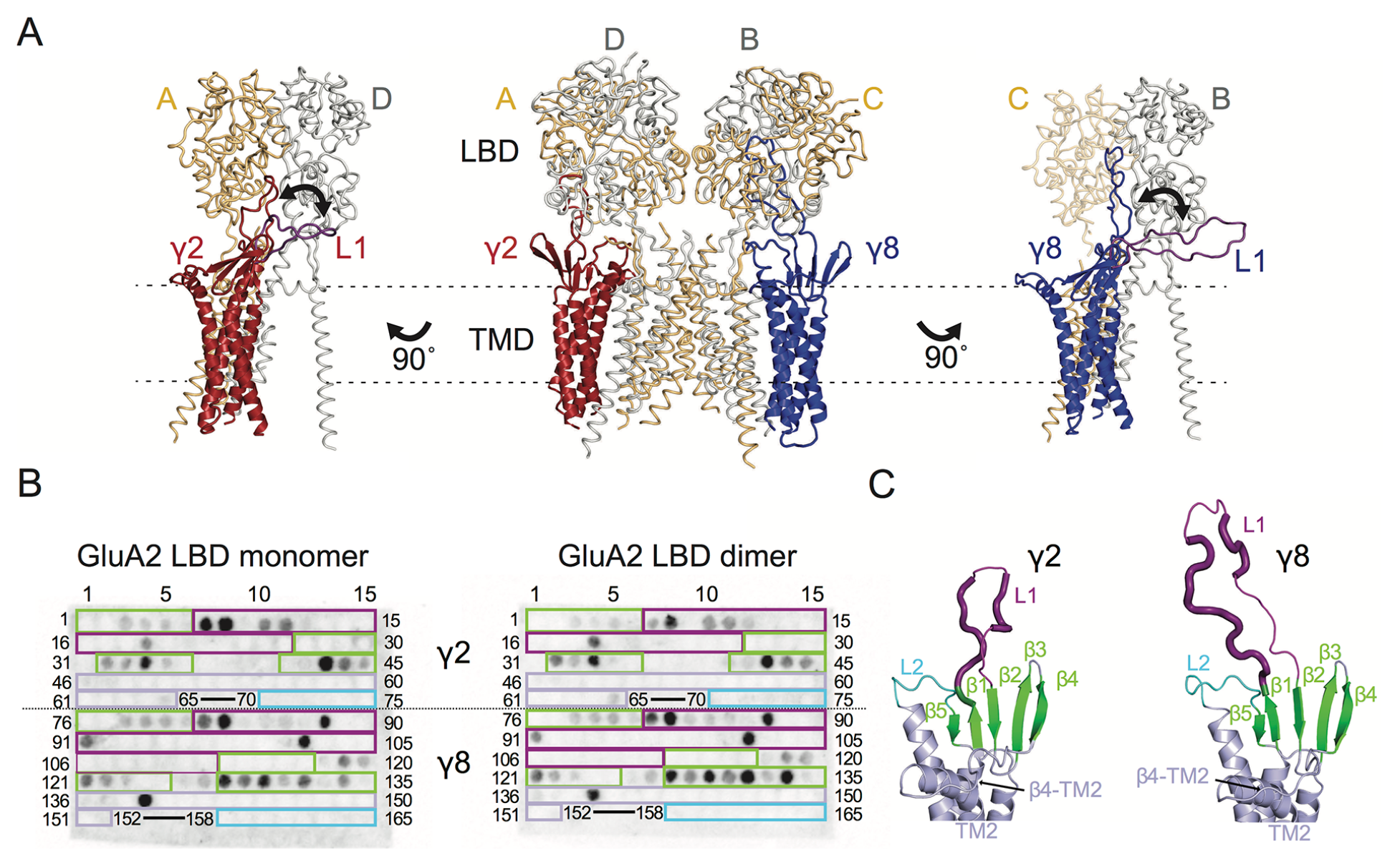
Modeling and biochemical analysis of AMPA-TARP complexes. **A**) The middle panel shows TARPS γ2 *(red)* and γ8 *(blue)* positioned between equivalent receptor subunits (A&D and B&C) based on the cryo-EM complex structure (5kk2). The predicted flexible extracellular L1 of γ8 is longer than in γ2 enabling it to reach more extensive regions of the receptor. To account for its flexibility we modeled L1 either between the LBD dimer (colored like the respective TARP) or underneath the lower lobe of the LBD *(purple; left panel for γ2, right panel for γ8)*. L1 might engage in different interactions with the LBD depending if located next to the inter-dimeric LBD interface (between subunits A & B or A & D; see Figure 1 – Figure supplement 1A). **B**) TARP peptide spotted membranes incubated with either monomeric *(left panel)* or dimeric GluA2 LBD *(right panel)*. Interacting peptides give a dark spot on the membrane when developed. The colored boxes indicate where the peptides are located in the TARPs (from β1 to L2) which is further illustrated in the structural models of γ2 and γ8 in panel C and Figure 1 – Figure supplement 1C. Quantitation of the spot arrays is found in Figure 1 – Source Data 1. **C**) Close up view on the modeled extracellular region of γ2 *(left)* and γ8 *(right)*. Secondary structure elements are shown in cartoon representation with *β*-sheets colored green, extracellular loop 1 in purple and loop 2 in cyan. Positive peptide hits in L1 are indicated by thicker loop-representation.

Repeated peptide mapping array assays indicated no clear preference for either monomeric or dimeric GluA2 LBD. However, in accordance with our hypothesis the majority of the L1 of both γ2 and γ8 contain hits in the peptide mapping array, indicating direct interaction with the receptor LBD (Figures 1B and C Figure 1 – Figure supplement 1C), albeit in conditions lacking the usual steric constraints of the complex. In the recent cryo-EM structures of the GluA2-TARP complex a possible interaction between a conserved negatively charged region located on the TARP β4-TM2 loop and the KGK motif in the lower lobe of the GluA2 LBD was predicted (34, 35). Thus we also tested for this potential interaction in the peptide mapping array but found no hits. A functional test of removing the acidic residues in this patch made γ2 into a much stronger modulator of AMPAR gating, with the steady-state current and superactivation both doubled (Figure 1 – Figure supplement 2). This result suggested that if interactions of the acidic patch with the receptor alter function, they actually inhibit the action of γ2. However, other sites have a dominant effect in the positive modulation of gating.

We also tested L2 of γ2 and γ8 for possible interactions with the LBDs because of its conserved charged features (4 and 7 charges), which are less prominent in γ5 and γ7 (3 and 1 charges respectively). Considering L2 being positioned distant underneath the LBD (around 15Å, measured between Cα of GluA2 P717 and γ2 K170 in the complex from PDB code: 5kbu (34) in the cryo-EM structures, it was not surprising that we found no interaction between L2 peptides and the GluA2 LBD. According to our GluA2-TARP models, in both γ2 and γ8 L2 is positioned between the S1-M1 and S2-M4 linkers (Figure 1 – Figure supplement 1A and B), which are outside the realms of our GluA2 LBD construct.

### Modulation of fast AMPA receptor gating by TARP L1 and L2 segments

To investigate the role of the extracellular domain of TARPs in controlling AMPA receptor activation, we made a series of chimeras and deletion mutants between γ2 and γ8. We first targeted the long loop in the first extracellular segment L1 (Figure 1) that has markedly different lengths and sequence content across the TARP family and its homologs. We also investigated the role of the shorter unstructured region in the second extracellular segment L2 (Figure 1), which is poised to interact with the LBD-TMD linkers of the AMPA receptor.

We first swapped L1 between γ2 and γ8 (Figures 2A and Figure 1 – Figure supplement 3), and assessed effects on desensitization. Although γ2 and γ8 apparently affect AMPA receptor desensitization similarly, γ8 slows down entry to desensitization more than γ2 (60 ± 5 s^−1^ and 40 ± 5 s^−1^, *n* = 24 and 9, for γ2 and γ8, respectively; Table 1). These chimeras exhibited asymmetric effects on desensitization. When activated by 10 mM glutamate, the chimera of γ2 with L1 from γ8 had steady-state current of 50 ± 5 % (*n* = 30; Figures 2A and D and Table 1), twice as large as γ2 alone (25 ± 2 %, *n* = 24 patches), and the rate of entry to desensitization was approximately halved (35 ± 5 s^−1^, *n* = 30; Figure 2C and Table 1). In contrast, the γ8 chimera with L1 from γ2 maintained the original desensitization behavior of the parent TARP (45 ± 1 s^−1^, *n* = 28; Figures 2A and C and Table 1). Deletion of L1 from γ2 and γ8 approximately halved the steady state current (15 ± 2 and 15 ± 3 %, *n* = 11 and 15, for γ2 ΔL1 and γ8 ΔL1, respectively; Figures 2B and D and Table 1), with a barely detectable speeding up of entry to desensitization (60 ± 5 s^−1^, *n* = 11 and 15, for γ2 ΔL1 and γ8 ΔL1, respectively; Figures 2B and C and Table 1). These results suggested that L1 can influence desensitization of complexes, as shown recently for GSG1L (30) but the absence of a simple exchange in desensitization behavior suggested that this loop functions in concert with other modulatory elements.

**Figure 2.**
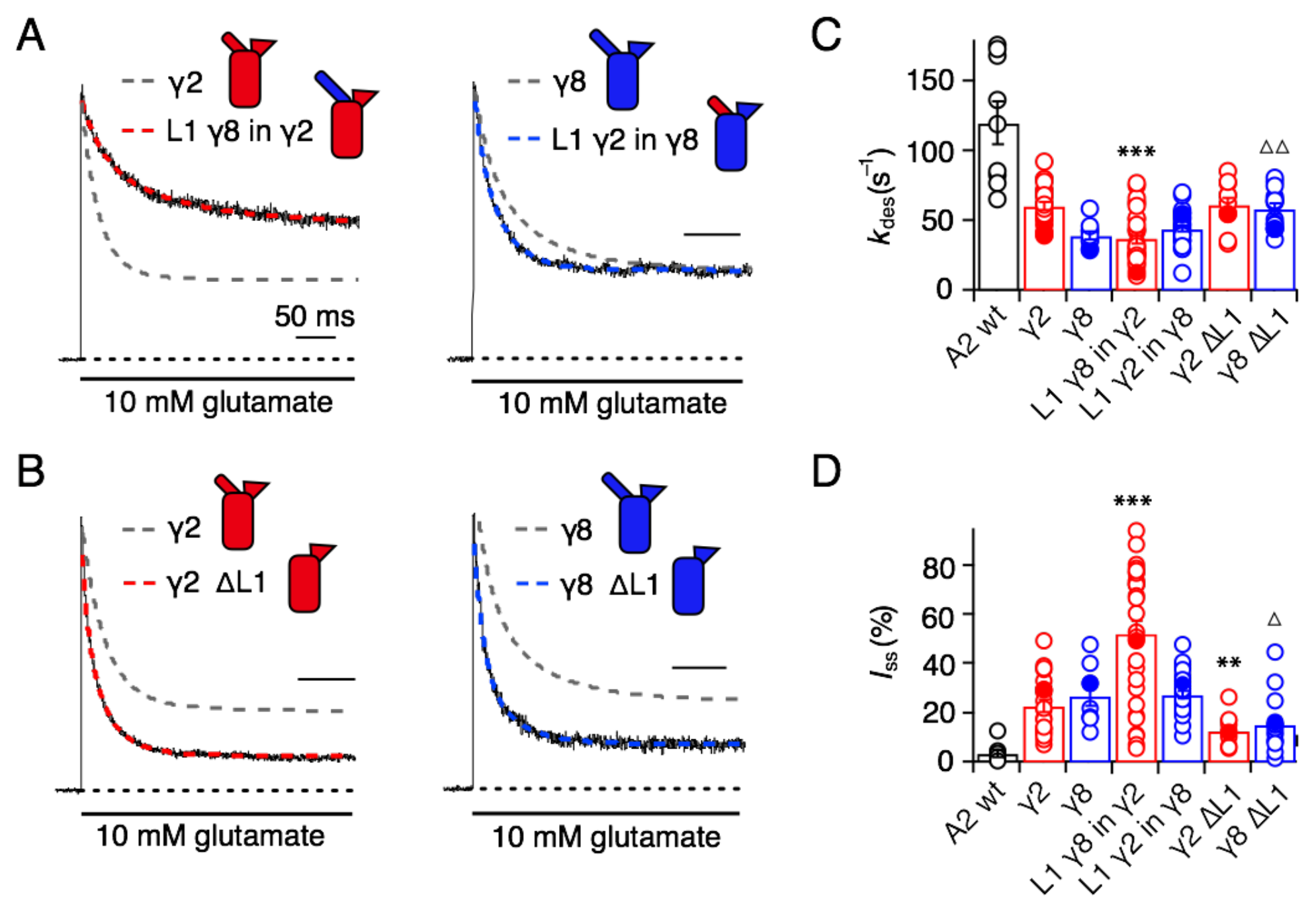
Desensitization properties of γ2 and γ8 L1 mutants. **A**) Representative traces from L1 γ8 in γ2 *(red)* and L1 γ2 in γ8 *(blue)* coexpressed with GluA2 in response to a 500 ms pulse of 10 mM Glutamate (*k*_des_ = 13 and 55 s^−1^; *I*_ss_ = 50 and 30 %, respectively). Example traces recorded from the parent TARPs coexpressed with GluA2 are shown in grey for comparison (*k*_des_ = 41 and 30 s^−1^; *I*_ss_ = 30 and 30 %, for γ2 and γ8, respectively). **B**) Representative traces from γ2 ΔL1 *(red)* and γ8 ΔL1 *(blue)* coexpressed with GluA2 in response to a 500 ms pulse of 10 mM Glutamate (*k*_des_ = 55 and 45 s^−1^; *I*_ss_ = 10 and 15 %, respectively). The wild type constructs coexpressed with GluA2 are shown as dashed lines for comparison. **C**) Bar graph summarizing the effects of the L1 mutation on the desensitization kinetics. **D**) Bar graph summarizing the effects of the loop1 mutations on the steady state current of the complexes. Currents were recorded at +50 mV in the presence of 50 *μ*M spermine in the pipette solution. For panels C and D, Filled symbols correspond to the traces shown in A) and B). \*\*\**p* < 0.001, ** *p* < 0.01, against γ2; Δ *p* < 0.05, ΔΔ *p* < 0.01, against γ8. Source data for panels C & D is found in Table 1 – Source Data 1. Error bars represent s.e.m.

**Table 1.**
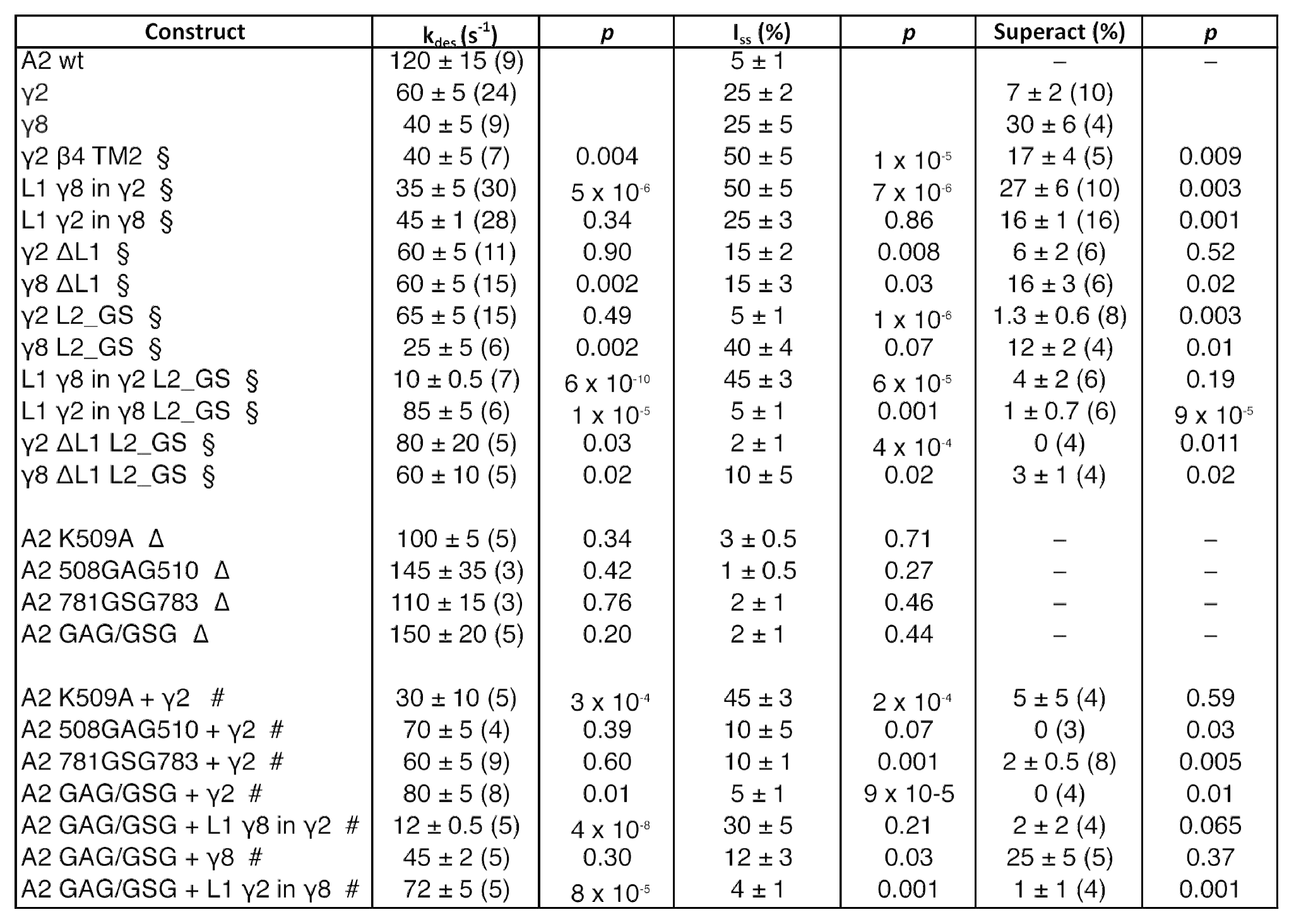
Kinetic properties of wild type and chimeric TARPs and GluA2 linker mutants. *k*_des_ is rate of desensitization, *I*_ss_ the steady state current expressed as percentage of the peak current and Superact the extent of superactivation expressed as the slow increase in steady state current during prolonged exposure to glutamate (see Materials and Methods for details). The number of patches recorded for each condition is shown in brackets. Values are shown as mean ± s.e.m. *p* values (from Student’s *t* test) are calculated as follows: § against the parent TARP; Δ against GluA2 WT; # against GluA2 WT + TARP. Currents recorded in the presence of TARPs were held at +50 mV in the presence of 50 μM spermine in the pipette solution. Recordings in the absence of TARPs were done at –60 mV without intracellular polyamines. Source data for Table 1 is found in Table 1 – Source Data 1

Seeking a further explanation for the modulation of desensitization by TARPs, we investigated the effects of altering the 8-residue stretch in the second extracellular segment of TARPs (L2), which connects TM3 to β5 in the extracellular domain. Replacement of the L2 segment with a flexible Gly-Ser linker, predicted to be of sufficient length not to disrupt the overall structure of the extracellular domain, had a striking effect on γ2. The rate of entry to desensitization was still slower than in receptors formed of GluA2 wild type (WT) alone (65 ± 5 s^−1^ and 120 ± 15 s^−1^, *n* = 15 and 9 patches for A2 + γ2 L2_GS and A2 WT, respectively; Figures 3A and C and Table 1), but the steady state current was reduced to the level of receptors without any TARP present (5 ± 1 % and 5 ± 1 %, *n* = 15 and 9 for A2 + γ2 L2_GS and A2 WT, respectively; Figures 3A and D and Table 1). In contrast, there was no detectable effect on γ8 of mutating this loop, except for a further slowing down of the desensitization rate (*k*_des_ = 25 ± 5 s^−1^, *I*_ss_ 4s = 40 ± 4%, *n* = 6, for γ8 L2_GS; Figures 3A, C and D and Table 1).

**Figure 3.**
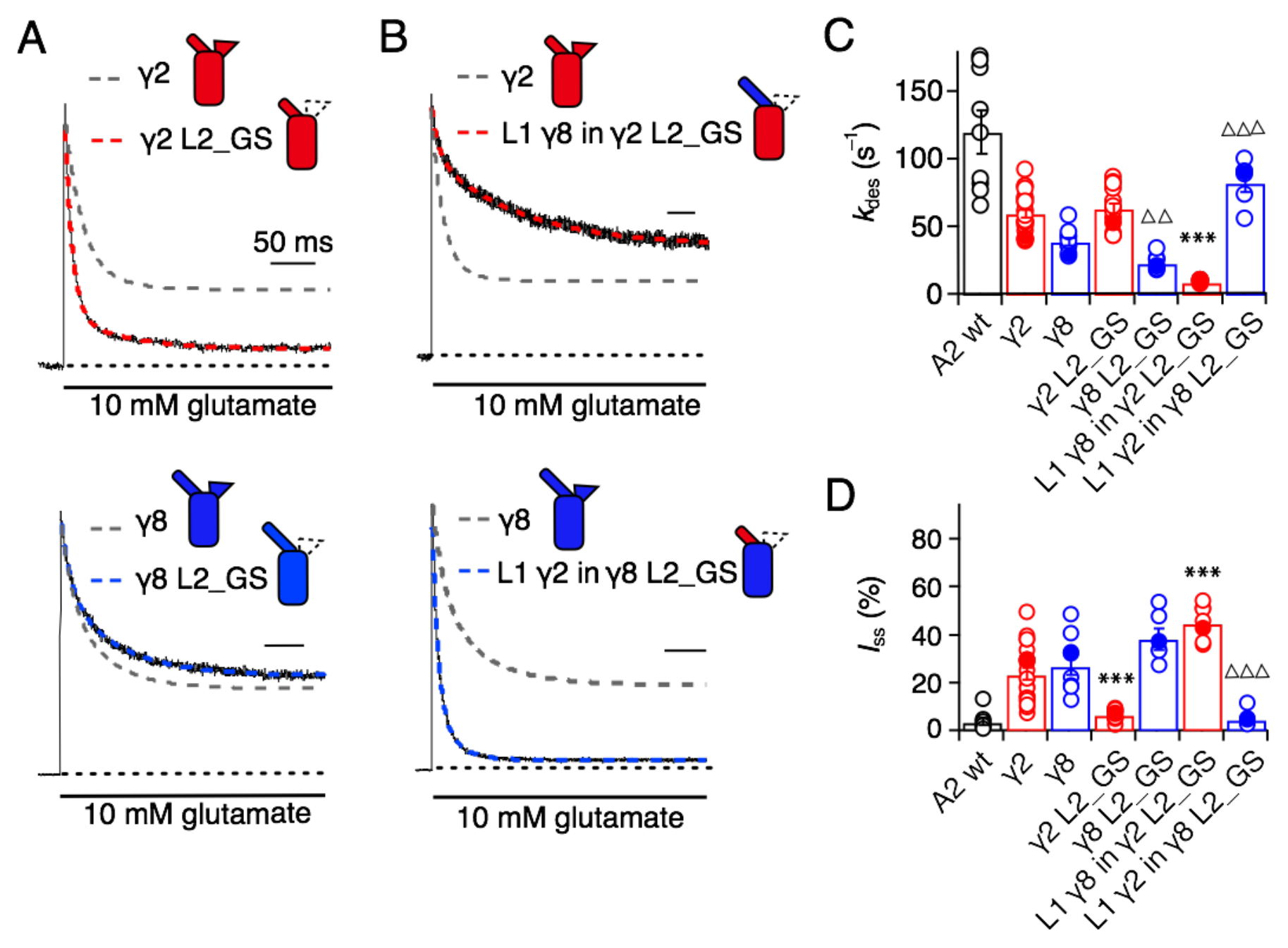
Desensitization properties of γ2 and γ8 L2 mutants. **A**) Neutralization of L2 in γ2 (γ2 L2_GS, *red*) decreased Iss, with little effect on γ8 (γ8 L2_GS, *blue*) (*k*_des_ = 50 and 20 s^−1^; *I*_ss_ = 5 and 35 %, respectively). Representative traces recorded from the parent TARPs are shown as dashed grey lines for comparison. **B**) Representative traces from L1 γ8 in γ2 L2_GS *(red)* and L1 γ2 in γ8 L2_GS *(blue)* coexpressed with GluA2 in response to a 500 ms pulse of 10 mM Glutamate (*k*_des_ = 10 and 90 s^−1^; *I*_ss_ = 40% and 5%, respectively). Traces from wild type γ2 and γ8 coexpressed with GluA2 are shown in grey for comparison. **C**) Bar graph summarizing the effects of the L2 mutation on the desensitization kinetics. **D**) Bar graph of the effects of the L2 mutation on the steady state current of the complexes. Filled symbols correspond to the traces shown in A) and B). \*\*\**p* < 0.001, against γ2; ΔΔΔ *p* < 0.001, ΔΔ *p* < 0.1, against γ8. Source data for panels C & D is found in Table 1 – Source Data 1. Error bars represent s.e.m.

Even more striking were results of coexpression of a chimera of γ2 with the GS-linker replacing L2, but harboring the long L1 loop of γ8. This chimera massively slowed entry to desensitization, producing complexes about 10-fold slower than receptors without any TARP (*k*_des_ = 10 ± 0.5 s^−1^, *n* = 7; Figures 3B and C and Table 1), and increased the steady state current during a 500 ms pulse of glutamate (45 ± 3%, *n* = 7; Figures 3B and D and Table 1). Making the inverse chimera (L1 from γ2 in γ8, with the GS-linker replacing L2) effectively nullified the modulatory activity of γ8.

The steady-state current was the same magnitude as for receptors that did not have γ8 (5 ± 1%, *n* = 6; Figures 3B and D and Table 1), and the rate of entry to desensitization (85 ± 20 s^−1^, *n* = 6; Figures 3B and C and Table 1) was closer to that of wild-type GluA2 than for the γ2 L2_GS chimera (see Table 1).

Although we performed all measurements at +50 mV, isolating heavily TARPed receptors by selecting for complexes with strong relief of polyamine block, we were concerned that some of the effects that we saw (particularly reduced or absent modulation) could be due to an altered stoichiometry of complexes, perhaps due to poor chimera expression. To assess this possibility, we measured the G-V relations for all the chimeras and deletion mutants (Figure 2 – Figure supplement 1). Importantly, all mutants gave responses that were strongly reduced in rectification, indicating that complex formation was normal. Broadly, each chimera closely followed the polyamine relief induced by the parent TARP, with γ2 chimeras producing populations of receptors that exhibited a greater rectification index than those based on γ8 (Figure 2 – Figure supplement 1).

### Superactivation of AMPA-TARP complexes

TARPs induce a subtype-specific superactivation of the GluA2 homomeric receptor. γ8 is a much stronger modifier of this slow gating mode than γ2 (36, 37). We investigated the role of the extracellular domain in superactivation using the same set of TARP mutants, but using 7-second applications of glutamate to measure the equilibrium level reached following superactivation. Our hypothesis was that the difference in superactivation between γ2 and γ8 would be specified by the sequence element most divergent between these two TARPs, L1.

In the chimeras swapping loop 1 between γ8 and γ2, the results were asymmetric (Figure 4). That is, loop 1 from γ8 could transfer the same degree of superactivation to γ2 (L1 γ8 in γ2, 27 ± 6 %, *n* = 10; Figures 4A and C and Table 1) but the reverse swap could not reduce superactivation to the level of γ2 (L1 γ2 in γ8, 16 ± 1 %, *n* = 16; Figures 4B and C and Table 1). The reason for this asymmetry became clear when we recorded complexes from which we removed L1 altogether from each TARP residual superactivation of 6 ± 2 and 16 ± 3% (for γ2 and γ8, respectively, *n* = 6; Table 1) were still present in the absence of L1. Therefore, although loop 1 can contribute to superactivation, and increase it over baseline levels, it is not the only element of TARPs driving this effect.

**Figure 4.**
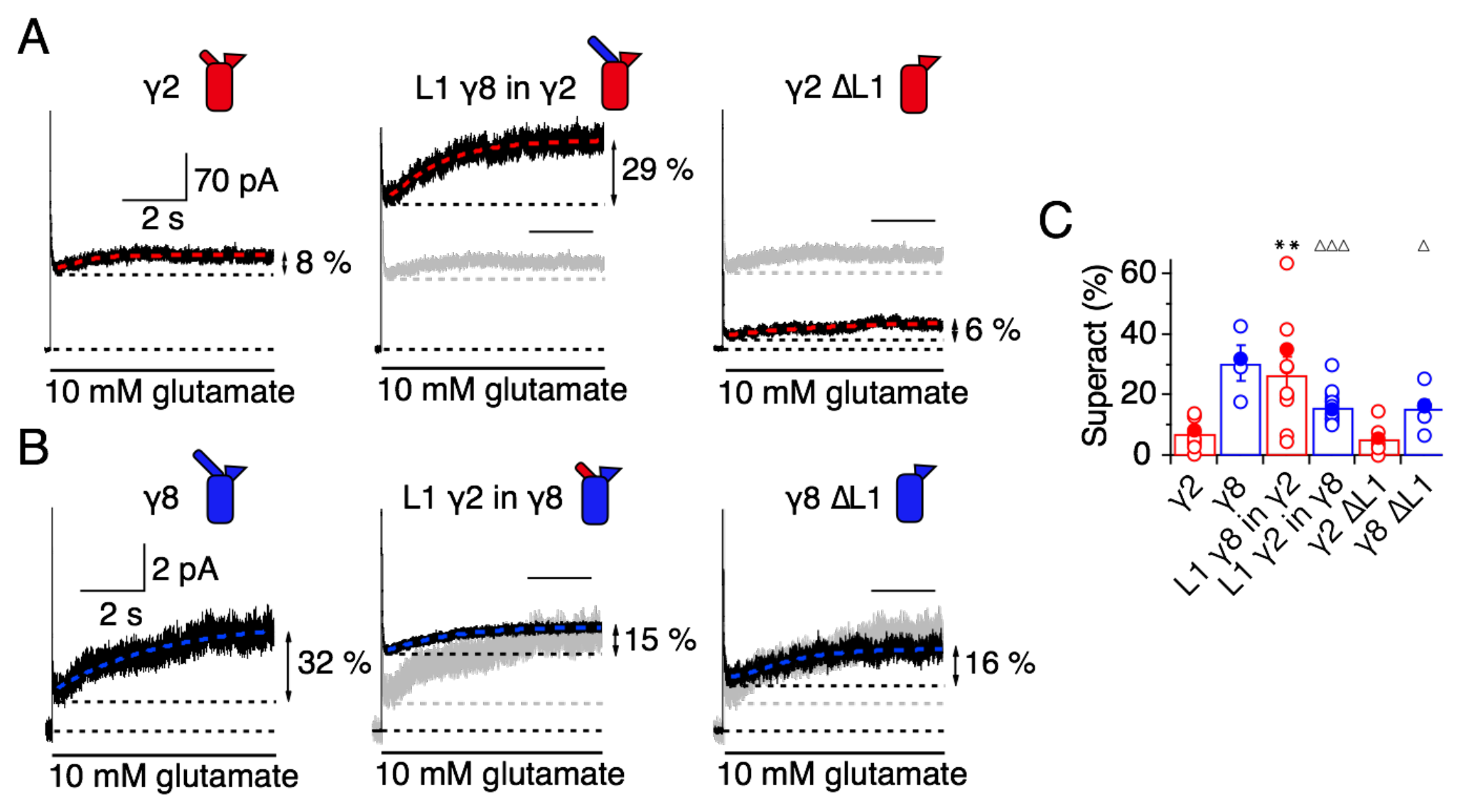
L1 modulates the extent of TARP-mediated superactivation. **A)** Example traces of γ2 wild-type and L1 mutants in response to 7 sec application of 10 mM glutamate. During prolonged application of 10 mM Glutamate γ2 induced superactivation of GluA2 receptors, shown as an increase in the steady state current (8% in the example shown, *left panel)*. The extent of superactivation was increased by 3-fold when L1 was replaced with that of γ8 *(central panel)*. Removing loop1 in γ2 did not affect superactivation much *(right panel)*. **B**) γ8 showed much bigger superactivation than γ2 during long glutamate exposure *(left panel)*. Shortening loop 1 by replacing it with that of γ2 or removing it decreased superactivation by 2-fold *(central and right panel)*. **C**) Bar graph summarizing the effects of the loop1 mutations on receptor superactivation. Currents were recorded at +50 mV in the presence of 50 *μ*M spermine in the pipette solution. Filled symbols correspond to the traces shown in A) and B) ** *p* < 0.01, against γ2; ΔΔΔ *p* < 0.001, Δ *p* < 0.05, against γ8. Source data for panel C is found in Table 1 – Source Data 1. Error bars represent s.e.m.

Given the residual superactivation that we saw in the absence of loop 1, we reasoned that loop 2 could play a role in receptor superactivation (Figure 5). We measured responses to 10 mM glutamate for the L2_GS mutants of γ2 and γ8 and found substantially reduced superactivation (1.3 ± 0.6 and 12 ± 2 %, *n* = 8 and 4, respectively; Table 1).

**Figure 5.**
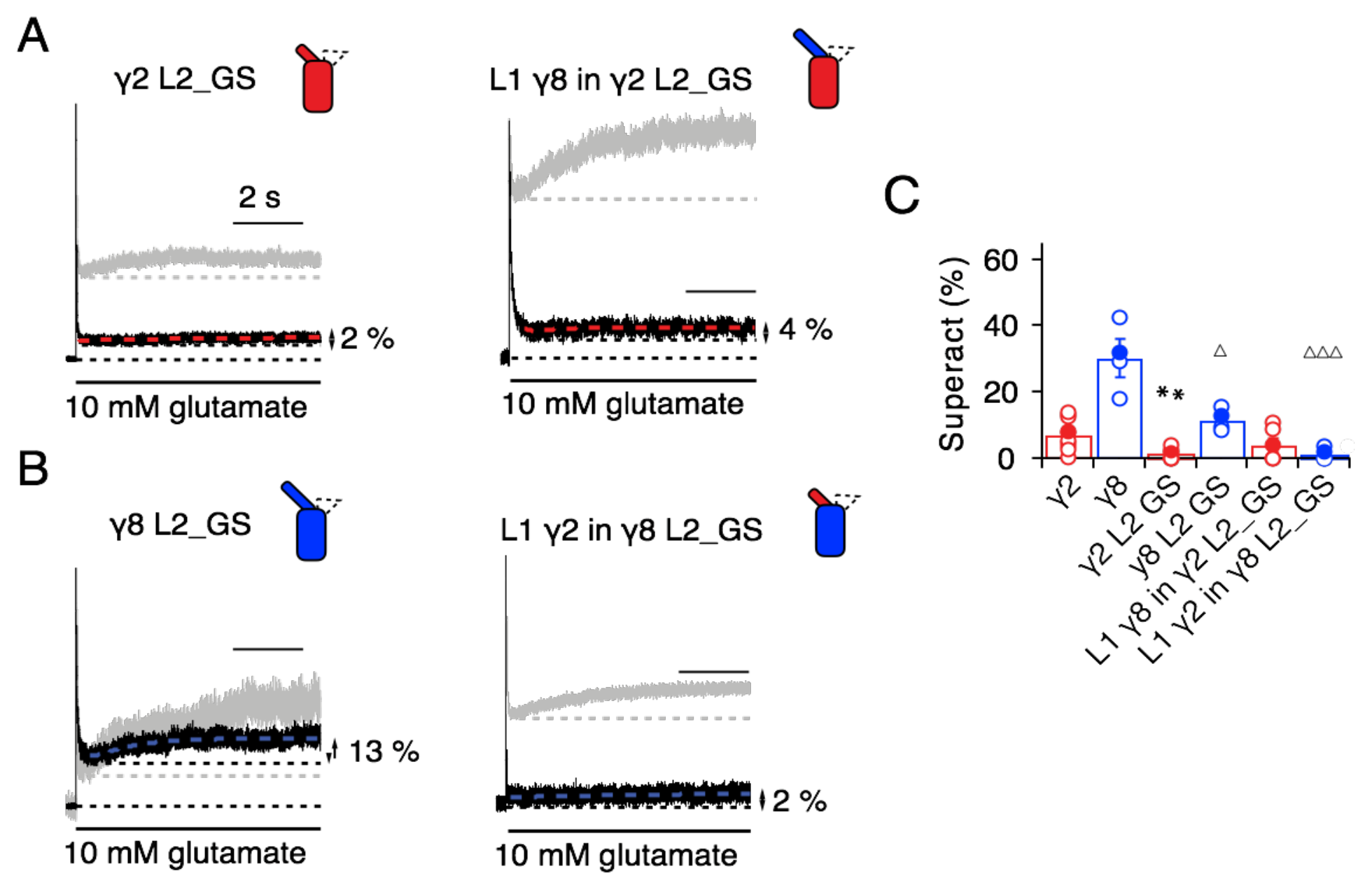
Superactivation of γ2 and γ8 L2 mutants. **A**) Neutralizing L2 from γ2 strongly reduced γ2-mediated superactivation *(left panel)*. On this background, L1 from γ8 induced only minimal superactivation *(right panel)*. They grey traces represent WT γ2 *(left)* and L1 γ8 in γ2 *(right)*. **B**) Removing L2 in γ8 decreased superactivation 2.5 fold *(left panel)*. Introducing L1 from γ2 on this background practically abolished superactivation *(right)*. The grey traces represent WT γ8 *(left)* and L1 γ2 in γ8 *(right)*. **C**) Bar graph of the effects of the L2 neutralization and L1 chimeras on superactivation. Filled symbols correspond to the traces shown in A) and B). ** *p* < 0.01, against γ2; ΔΔΔ *p* < 0.001, Δ *p* < 0.05, against γ8. Source data for panel C is found in Table 1 – Source Data 1. Error bars represent s.e.m.

Even more strikingly, the same TARP mutants with loop 1 swapped had a further reduced effect. The loop 1 from γ2 in the L2_GS mutant of γ8 had almost negligible superactivation, reduced by ~15-fold from wild-type γ8, to about 1 ± 0.7 % (*n* = 6; Figures 5B and C and Table 1). Taking into account the lack of steady-state current, fast desensitization and similar deactivation kinetics to wild-type GluA2 alone that we observed in patches containing complexes of GluA2 with the L1 γ2 in γ8 L2_GS mutant, we classed this chimera as a kinetic null of γ8.

The TARP chimeras that exhibited the least power to slow desensitization kinetics and to stabilize active states were those that replaced charged residues in the L2 segment, and from which we either deleted L1, or included the short loop from γ2. These observations guided our construction of a kinetically-null γ2. We reasoned that a γ2 chimera lacking L1 and with a GS-linker replacing L2 should associate normally with GluA2 but might have no kinetic effect at all on the receptor complexes. Indeed, γ2 ΔL1 L2_GS associated normally into the receptor complex (as assessed by relief of polyamine block, Figures 6A and B) but this mutant γ2 was highly deficient in modulating gating of GluA2. Superactivation, and the increase in steady state current were absent in these complexes (superactivation = 0 %; *I*_ss_ = 2 ± 1 %, *n* = 4 and 5, respectively; Figures 6C and D and Table 1). Somewhat surprisingly, the deletion of L1 from γ8 on the L2-GS background retained a larger steady state current than the chimera that included the L1 segment of γ2 (*I*_ss_ = 5 ± 1% and 10 ± 5 %, *n* = 6 and 5, for L1 γ2 in γ8 L2_GS and γ8 ΔL1 L2_GS, respectively; Figures 5B and C, 6E and G and Table 1) and a small superactivation (3 ± 1 %, *n* = 4; Figure 6A and Table 1).

**Figure 6.**
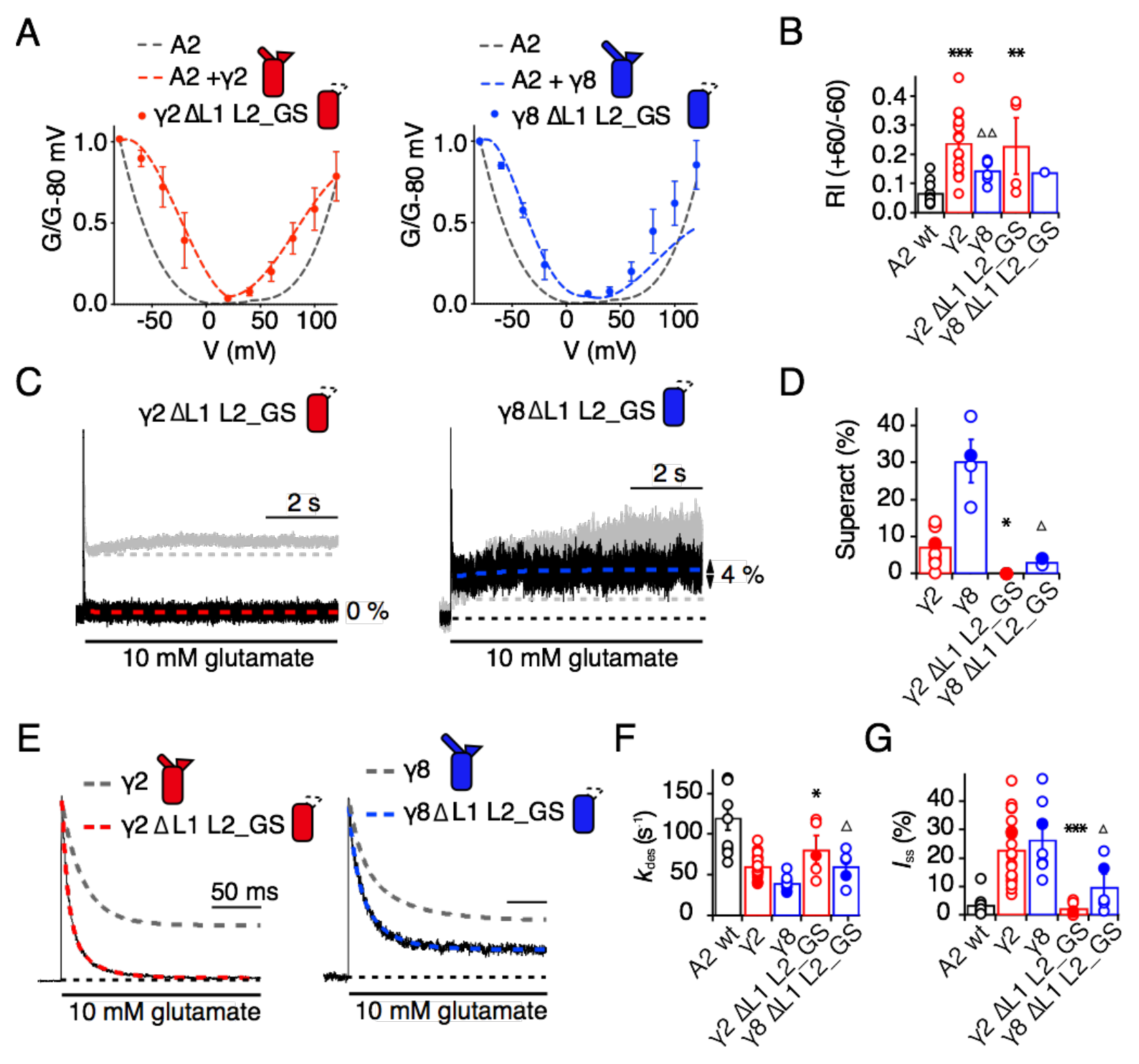
Eliminating L1 and L2 removes modulation by γ2. **A)** Mutation of both L1 and L2 in γ2 *(left panel, red)* and γ8 *(right, blue)* did not change association of TARPs with AMPA receptors, as assessed by the G-V curve. GluA2 WT is shown in grey. **B**) Bar graph summarizing the rectification index of the dual loop mutations. **C**) Example traces of γ2 ΔL1 L2_GS *(left)* and γ8 ΔL1 L2_GS *(right)* in response to 7 sec application of 10 mM glutamate. Corresponding wild-type TARPs are shown as dashed lines. **D**) Bar graphs summarizing the effects of the dual loop mutation in γ2 *(red)* and γ8 *(blue)* on superactivation. **E**) Representative traces from γ2 ΔL1 L2_GS *(left)* and γ8 ΔL1 L2_GS *(right)* coexpressed with GluA2 in response to a 500 ms pulse of 10 mM Glutamate (*k*_des_ = 74 and 50 s^−1^ *I*_ss_ = 1.5 and 16 %, respectively). Currents from the parent TARPs are shown in grey for comparison. **F**) Bar graphs summarizing the effects of the dual loop mutation in γ2 *(red)* and γ8 *(blue)* on desensitization decay. **G**) Bar graph summarizing the effects of the double loop mutation on the steady state current of the complexes. Currents were recorded at +50 mV in the presence of 50 *μ*M spermine in the pipette solution. For panels D, F and G, filled symbols correspond to the traces shown in C) and E). \*\*\**p* < 0.001, ** *p* < 0.01, * *p* < 0.05, against γ2. Source data for panel B is found in Figure 6 – Source data 1. Source data for panels D, F & G is found in Table 1 – Source Data 1. Error bars represent s.e.m.

### L2 controls gating through interaction with linkers proximal to the channel gate

From our models, a range of sites on GluA2 could interact with L1, including the KGK motif in the LBD (30, 38). Substitutions at L2 of γ2 and γ8 had profound effects on gating of TARP complexes and are well placed to interact with gating machinery (Figure 1A and S1B). Particularly, we expected from our structural models and other available structural data (34, 35) that L2 should interact with the S1-M1 linker and the S2-M4 linker in the AMPA receptor. The L2 sequence has an alternating charge motif that is mirrored in two parts of the GluA2 linkers 508–510 and 781–783. These segments are immediately adjacent to the TARP L2 in all four subunits.

Replacement of 508QKS510 to GAG in the S1-M1 linker (GluA2 508GAG510, Figure 7A) produced a GluA2 receptor with normal kinetics and that associated normally with γ2 and γ8 (Figure 7 – Figure supplement 1). Strikingly, in complexes with WT γ2, this mutant phenocopied the neutralizing truncation of L2 in TARPs well (see Figure 3), abolishing superactivation and reducing the steady state current (0% and 10 ± 5%, *n* = 3 and 4, for superactivation and *I*_ss_, respectively; Figures 7C-E and Table 1). In contrast, a point mutant K509A, also with normal gating (Figure 7 – Figure supplement 1), was more strongly modulated by γ2, providing further indication that a second site was potentially involved (Figure 7E and Table 1). Our model suggested that the S2-M4 linker of GluA2 was equally well positioned to interact with L2 from γ2. To test the importance of the alternating charges in the S2-M4 linker, we made another triple mutation replacing 781KEK783 to GSG (GluA2 781GSG783, Figure 7B). This mutant again had normal kinetics in the absence of γ2 (Figure 7 – Figure supplement 1), but also exhibited a reduced steady state current and negligible superactivation (10 ± 1 % and 2 ± 0.5%, *n* = 9 and 8, for Iss and superactivation respectively; Figures 7C-E and Table 1). Importantly, the combination of these two triple mutants, abolished the entire modulatory effect of γ2 on the AMPA receptor, reducing superactivation and the instantaneous steady-state current to the same level as GluA2 in the absence of TARP (0 % and 5 ± 1 %, *n* = 4 and 8, for superactivation and *I*_ss_, respectively; Figures 7C-E and Table 1). This mutant receptor retained ostensibly normal gating and association to TARPs (Figure 7 – Figure supplement 1), despite the absence of gating modulation.

**Figure 7.**
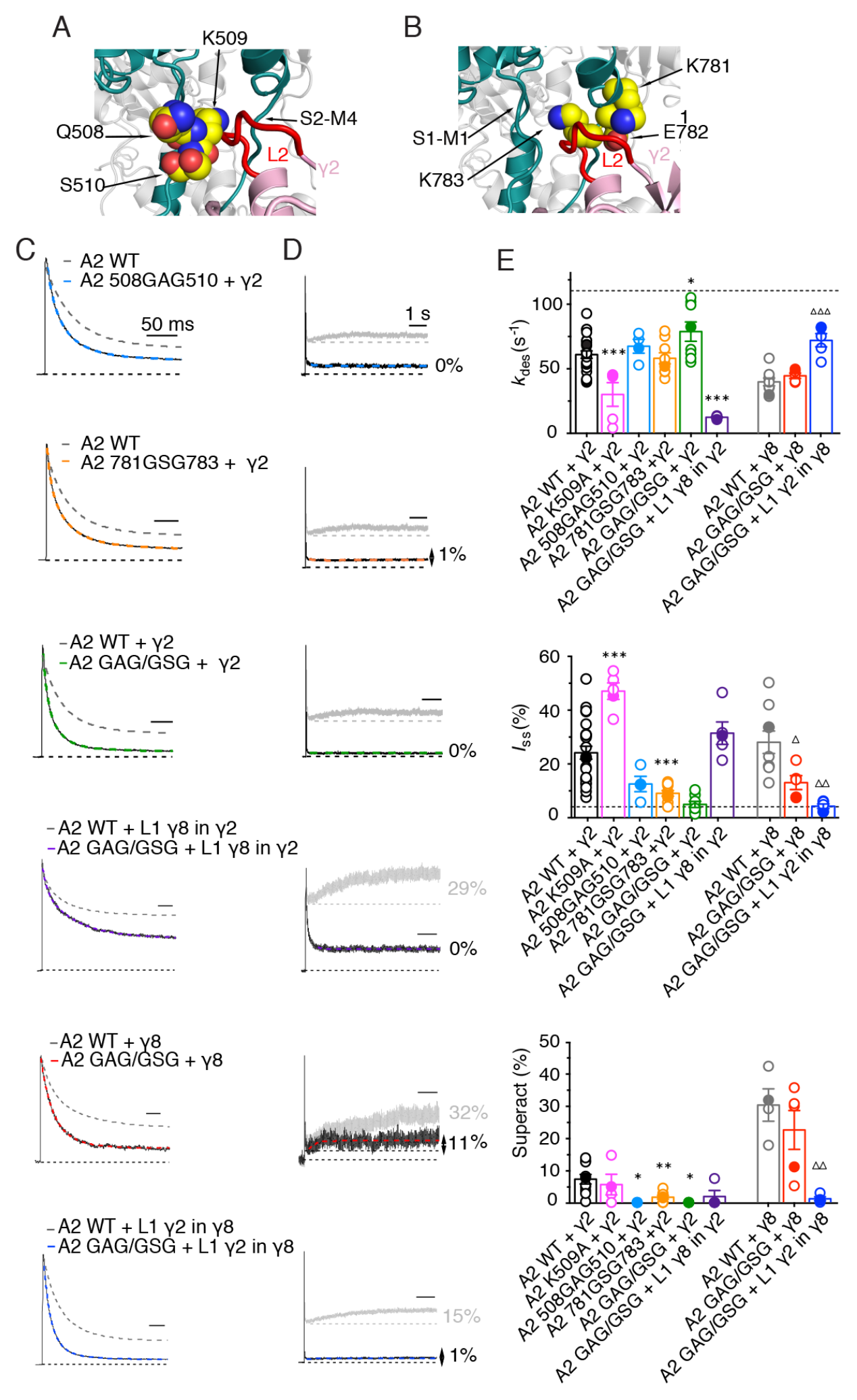
The LBD-TMD linkers are the key sites for modulation of AMPA receptor gating by TARPs. **A)** Residues in the S1-M1 linker (Gln508, Ser509, and Lys510 represented as yellow atomic spheres) are in close proximity to the L2 of TARPs (L2 of γ2 is shown in red). **B**) Residues in the S2-M4 linker (Lys781, Glu782 and Lys783) predicted to interact with L2 are labeled and shown as yellow atomic spheres. **C**) Example responses from linker mutants coexpressed with γ2, γ8 and loop 1 chimeras to 500 ms 10 mM Glutamate. D) Representative responses from linker mutants coexpressed with γ2, γ8 and loop 1 chimeras to a 7 sec pulse of 10 mM Glutamate. The extent of superactivation is indicated. **E**) Bar graphs summarizing the desensitization properties *(top panel)*, steady state current *(central)* and superactivation *(bottom)*. Colors are as in panel C. Filled symbols correspond to the traces shown in panels C and D. \*\*\**p* < 0.001, ** *p* < 0.01, * *p* < 0.05, against γ2; ΔΔΔ *p* < 0.001, Δ *p* < 0.05, against γ8. Source data for panel E is found in Table 1 – Source Data 1. Error bars represent s.e.m.

To discern whether the loss of modulation occurred because the linker sites are the primary interaction site, or whether the linkers both interact with TARPs and transmit upstream modulation from sites in the LBD, we assessed modulation by γ8 and related chimeras. The propensity of γ8 to modulate gating of the double linker mutant (GluA2 GAG/GSG) was reduced, but robust superactivation could still be observed (25 ± 5%, *n* = 5, Figure 7 and Table 1). Given this result, which suggested that L1 could still modulate gating of complexes, we hypothesized that the γ2 chimera incorporating the L1 of γ8 should also modulate the double linker mutant. This chimera could not produce superactivating complexes (2 ± 2 %, *n* = 4, Figures 7D and E, as for the γ2 chimera lacking L2 interactions, L1 γ8 in γ2 L2_GS, Figure 5A) but retained the slow desensitizing behavior due to L1 (*k*_des_ = 12 ± 0.5, *n* = 5, Figure 7E and Table 1).

In coherence with our previous results, mutation of the GluA2 linkers ablated the effect of the γ8 chimera with L1 from γ2 to modulate the kinetics of complexes, reducing the steady state current and superactivation to the same levels as GluA2 wild-type in the absence of TARP (*I*_ss_ = 4 ± 1 %, superactivation = 1 ± 1 %, *n* = 5 and 4, Figure 7E and Table 1). Therefore, in the absence of the long L1, γ8 fails to modulate GluA2 when the S1-M1 and S2-M4 linker interaction sites are removed (again consistent with its cousin lacking L2 interaction sites, the L1 γ2 in γ8 L2_GS variant; see Figure 5C).

Overall, these results indicate that the long loop of γ8 L1 is still able to modulate complexes at extracellular sites with the receptor linker sites disrupted, supporting the idea that the linkers do not function primarily to transduce distant TARP modulation. Rather, the LBD-TMD linkers are the primary modulatory site for both γ8 and γ2. The latter has a short L1 loop, and cannot modulate receptors if the L2 interaction is absent. However, γ8 combines the longer L1 and the L2 site to modulate receptor properties more effectively, in a compound fashion.

## Discussion

The results we present here offer several new insights into TARP function. First of all, extracellular sites account for all the modification of AMPA receptor gating by TARPs. Previous work showed that L1 could transfer aspects of modulation between TARPs, but our experiments indicate that the 2nd short extracellular segment (L2), which varies strongly in sequence between TARPs, is dominant. Further work will be required to establish the generality of this modulatory mechanism.

Secondly, these same sites do not have any appreciable role in determining assembly of TARP-AMPA receptor complexes. Intuitively, this division of roles makes sense because gating modification requires transient interactions on a timescale far faster than receptor assembly. Therefore, interactions between transmembrane segments and intracellular regions are responsible for assembly and modulation of polyamine block.

Thirdly, we show that the linkers to the transmembrane domain are key sites for modulation of AMPA receptor gating by auxiliary proteins, and provide insights into the molecular basis of this interaction. Previous work suggested ATD interactions and prominent roles for the LBD in modulation, but the interactions we demonstrate here are much more proximal to the channel gate (25). We could show a very close functional confluence between modifying the receptor itself and modifying each TARP, at an interaction site predicted from structural modeling. The elimination of modulation by nullifying L2 of γ2, or by mutating residues in the LBD-TMD linkers of GluA2, strongly implicates this site as a pivotal interaction underlying modulation. Putative electrostatic interactions posited from structural studies require a large conformational change (between 13 Å and 25 Å depending on the TARP’s position in the complex; measured between C-alpha atoms from GluA2 K699 and γ2 D92 in cryo-EM complexes 5kbu and 5kk2, respectively) (34, 35). A key point here is that these interactions are secondary to those involving L2 at the AMPAR linkers. These interactions should occur readily for each auxiliary protein subunit, allowing a maximal 4:4 stoichiometry with minimal conformational change for γ2 (Figure 8A) (35). For other auxiliary proteins, for example γ8, the stoichiometry of the L2-linker interaction would vary with the number of associated TARPs, but will not be limited by position of the TARP within the complex (Figure 8B). Finally, neutralization of the major part of the acidic patch strongly enhanced modulation of gating by γ2, ruling out that negative charges here have a dominant role in modulation.

**Figure 8.**
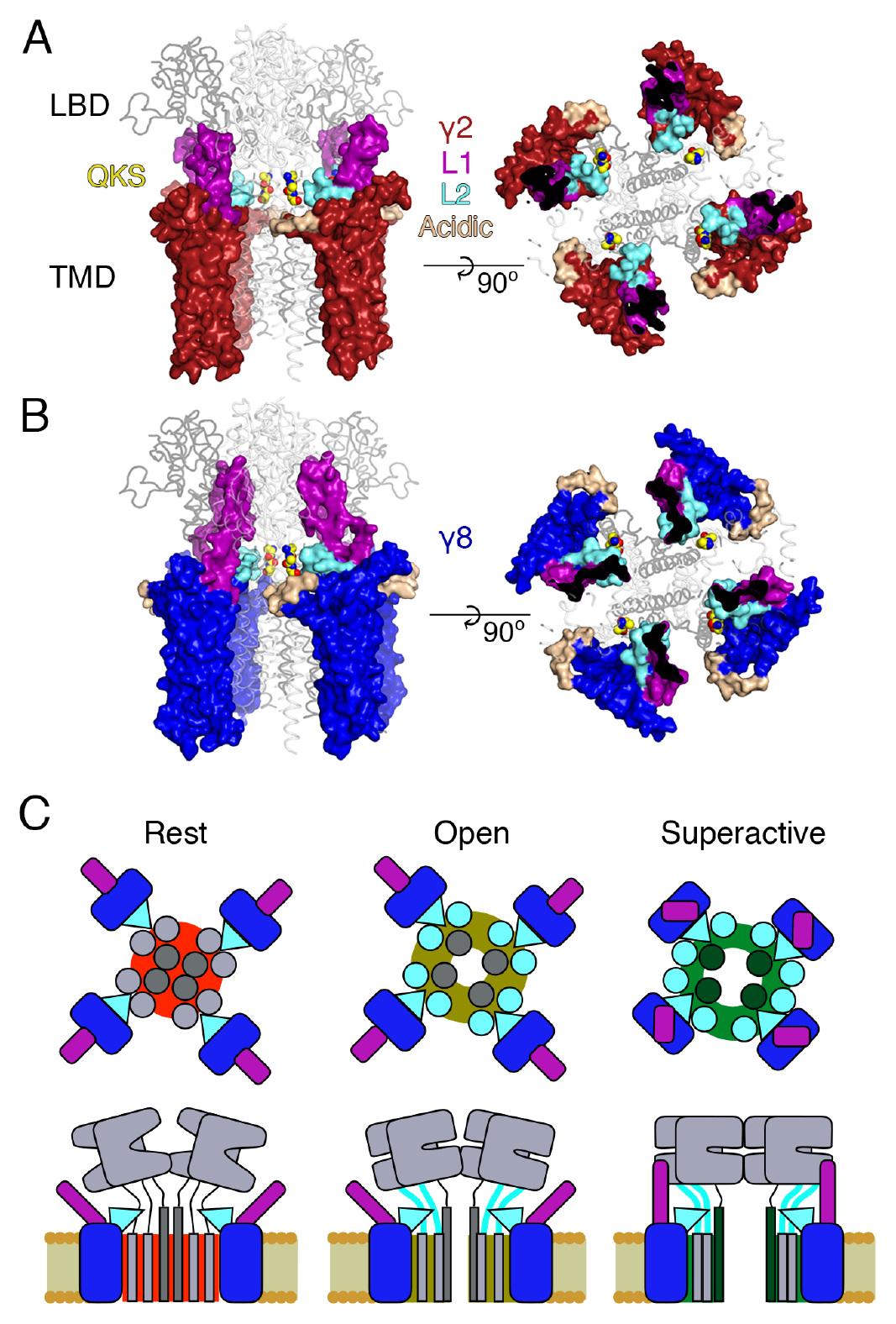
Proposed mechanism of AMPA modulation by TARPs. **A** Model of a AMPA-γ2 complex in front view *(left)* and top view *(right)*. Four molecules of γ2 *(red)* are shown with L1 and L2 colored in magenta and cyan, respectively. L2 is sandwiched between the LBD-TMD connecting linkers of the receptor (grey, amino terminal domains omitted for clarity). The QKS sequence on the S1-M1 linker is shown as yellow atomic spheres. The acidic patch on the β4-TM2 linker is indicated in wheat. **B**) The model of γ8 *(blue)* shows the similar interactions of L2 *(cyan)*. The orientation of the more extensive loop 1 of γ8 is not known, here it is depicted reaching up to the LBD dimer. **C**) Cartoon model of the proposed AMPA modulation mechanism, taking the example of γ8. The AMPA-TARP complex is shown from top (upper panel) and in side view (lower panel). The receptor is colored in grey (pore forming M3 domain depicted in dark grey). γ8 is colored as in panel B, with the acidic patch omitted. In the resting state (indicated by a red, closed pore) L2 is positioned in close proximity to the LBD-TMD connecting linkers. Once glutamate binds to the LBD, the resulting conformational change is transduced via the LBD-TMD linkers to open the pore (olive green, open state). During this transition L2 could wedge between the S1-M1 and S2-M4 linkers to modulate the receptor gating. The concerted action of L1 and L2 is necessary for superactivation of the receptor (dark green, high open probability state), most likely via L1 to stabilizing the LBDs layer.

Fourth, we show that the long extracellular loop 1 of γ8 is a very strong positive modulator of AMPA receptor gating, whose influence is likely held in check by the substoichiometric combination of γ8 with the AMPA receptor (28). The subunit γ8 slows receptor desensitization via L1. This loop can produce a profound block of desensitization when transplanted to γ2, and probably interacts state-specifically with the LBD dimer because of its substantial reach (for examples see Figures 1 and 8). Previous kinetic measurements suggest that superactivation is adopted by a minor population of receptors in equilibrium with saturating glutamate, speaking in favor of a weak interaction that is boosted by the high effective concentration of L1 close to its site of action in the receptor complex.

Our approach to fit Claudins with modeled loops from TARPs into the best resolution cryoEM reconstruction available (5KK2, (35)) has clear implications for modulation. Our model, when compared to the independently derived model of TARP-AMPA modulation (34), presents the TARPs oriented at a subtly different angle. Therefore, our model predicted the L2 interaction on the basis of one set of CryoEM data. We could not adequately incorporate the loops and the original structures of the receptor linkers in this model (Figures 1 and S1). Whilst this problem could be due to deficits in our model, another explanation is that the linkers (S1-M1 and S2-M4) are disrupted from their basal positions, and that the L2 loop can wedge between them. Upon activation, it is expected that the linkers will move away from the overall pore axis, which could permit further state-dependent interactions (See cartoon in Figure 8C).

Future structural studies may permit a more detailed view into the interactions between L2 and the linker domains of AMPAR. Although Claudin structures allowed positioning of auxiliary proteins with high confidence within CryoEM reconstructions, the loops that we have investigated here are not resolved within these structures, possibly because they interact transiently and are otherwise disordered. Although our peptide array suggested that stretches of L1 interact with the LBD, we were not able to obtain co-crystal structures of peptides with monomeric or dimeric forms of the GluA2 LBD. Nonetheless, knowledge of Claudin structures enabled us to make structurally sympathetic substitutions into TARPs for functional experiments that did not disrupt expression or assembly of complexes. These approaches are in contrast with most previous work which simply swapped extracellular portions, including mismatching the folded portions of the TARP extracellular domain. Two observations highlight the importance of sympathetic exchanges. First, some naive deletions would be expected to alter TARP structure. The simple deletion of L2 would severely disrupt the extracellular domain of γ2 or γ8, because this segment connects structured regions separated by about 10 Å. Second, some deletion chimeras we made retained modulation, with the most striking example being γ8 ΔL1 L2_GS, which retained a substantial steady state current (Figure 6). The residual modulation could be related to the presence of a few residues from L1 in the γ8 ΔL1 L2_GS (see Figure 1 – Figure supplement 3). Without maintaining these residues, the chimera did not express. This observation illustrates the sensitivity of domain boundaries in TARPs.

Because our observations suggest that the AMPA receptor linkers are key to TARP modulation, it is likely that chimeric receptors with altered linkers that exhibit constitutive gating are bad reporters of the TARP-GluA modulation, although they clearly delineate assembly motifs (29). The molecular nature of the interactions we have identified here raise the intriguing possibility that acute disassembly of complexes, rather than modulation, might be the target of recent subtype specific drugs (19, 20).

Our results allow us to construct a tentative model for the distinct forms of modulation that TARPs produce (Figure 8C). The slow increase in glutamate efficacy, which we term superactivation, is specified by the combination of L1 and L2, whereas the basal increase in steady state current arises from L2 alone. We previously modeled the modulatory interaction between TARPs and the AMPA receptor with single conformational change, but did not consider desensitization. The concerted involvement of multiple loops suggests multiple conformational states are required to describe the interaction, most notably in the case of γ8. The greater conformational space that can be explored by loop 1, and its strong connection to superactivation, indicate that these conformational changes could relate to the slow transitions represented in the model of superactivation (37). In contrast, conformational changes of the linker region of the AMPA receptor upon opening will naturally lead to a state-dependent interaction with L2 of γ2 or γ8, because of the direct proximity. A further level of complexity is that an intact L2 segment is required for the strong superactivation induced by γ8, but is not required at all for slow desensitization behavior that the long L1 loop of γ8 can produce. Because in these experiments, slow desensitization occurs when occupancy of superactive states is low, we can quite reasonably assume that L1 adopts multiple conformations to stabilize separate functional states of the receptor, and that some functional signatures require a concerted action of both loops. Additional stabilization of desensitized states by the variable loop 1 is also likely (30).

This work has produced mutant TARPs and AMPA receptors that both lack modulatory properties, and also those that have greatly enhanced modulation. Both these signatures of activity should be useful tools for investigating TARP action in synapses, including understanding the relative importance of assembly into complexes for anchoring (39) as opposed to kinetic modulation, for clarifying the consequences of TARP modulation for short term plasticity (18), and for better identifying TARPs in ternary complexes with other auxiliary subunits (17, 40).

## Materials and Methods

### Molecular biology

We used GluA2 flip receptors, unedited at the pore site (Q-containing) in the pRK vector also expressing eGFP following an internal ribosomal entry site (IRES) sequence. Mouse γ2 was the kind gift of Susumu Tomita and was expressed from an IRES-dsRed construct as previously described (37). Mouse γ8 (the kind gift of Roger Nicoll) was expressed the same way. Point mutations and chimeras were created by overlap PCR and confirmed by double-stranded sequencing. The construct boundaries of the chimeras used are shown in Figure 1 – Figure supplement 3. Residues in GluA2 were numbered based on the assumption that the signal peptide is 21 residues.

### Patch clamp electrophysiology

Wild type or mutant GluA2 and TARP constructs were co-transfected in HEK 293 cells with PEI. The ratios of co-transfection were 1:2 for GluA2-γ2 and 1:5 for GluA2-γ8, up to 2 μg total DNA per 35 mm dish. The same ratios were maintained for all the reciprocal mutants. Cells were supplemented with 40 μM NBQX to reduce TARP-induced cytotoxicity. Recordings were performed 24–48 hours after transfection. The external recording solution contained (in mM): 150 NaCl, 0.1 MgCl_2_, 0.1 CaCl_2_ and 5 HEPES, titrated to pH 7.3 with NaOH. The pipette solution contained (in mM): 120 NaCl, 10 NaF, 0.5 CaCl_2_, 5 Na_4_BAPTA, 5 HEPES and 0.05 spermine, pH 7.3. 10 mM glutamate was applied to outside-out patches with a piezo-driven fast perfusion system (PI, Germany). In order to isolate currents exclusively mediated by TARPed receptors, patches were voltage-clamped at a holding potential of +50 mV. Currents were low-pass filtered at 5 kHz using an Axopatch 200B amplifier (Molecular Devices, U.S.A.) and acquired with Axograph X software (Axograph Scientific, U.S.A.). Typical 10–90% solution exchange times were faster than 300 μs, as measured from junction potentials at the open tip of the patch pipette.

#### Data analysis

To measure receptor desensitization we applied 10 mM glutamate for 500 ms. Desensitization rate and steady-state current were then obtained by fitting the traces with a sum of two, and when necessary three, exponentials. Rates constants are expressed as weighted mean of multiple components. Superactivation was measured during a 7 second application of glutamate and was defined as the excess steady-state amplitude following the desensitization trough, normalized to the peak current. A triple exponential function was used to fit the slowly augmenting current of superactivation measurements. To account for possible variability in the response and expression of the complexes, we tried to record at least 5–6 patches from at least three different transfections for each condition. For experiments with very low success rates (that is, worse than 1 patch in 20 giving an acceptable recording), in the presence of γ8, at least three patches were collected. No data were excluded, except from patches where recordings were unstable, had excessive rundown or solution exchange slower than 0.5 ms as measured after the experiment. Results are shown as mean ± standard error of the mean (s.e.m.) and statistical significance was assessed with a two-tailed Student’s *t*-test as specified in Table 1.

### Protein expression and purification of soluble LBDs

Using the flop isoform of rat GluA2 ligand binding domain (S1S2 fusion) in pET22b vector (kindly provided by E. Gouaux) as a base, we inserted the flip mutations N744T, A745P, N754S, L758V, and added the C-terminal residues Lys776-Gly779 (GluA2_LBD) and the non-desensitizing mutation L483Y (GluA2_LBD_LY) by overlap mutagenesis. Protein expression and purification was carried out as described previously (41). Briefly, monomeric and dimeric (L483Y) LBDs were expressed in *E. coli* Origami B (DE3). Cells were harvested by centrifugation, lysed and subjected to metal affinity chromatography and size exclusion chromatography. Fractions containing the N-terminal His_8_-tagged protein were pooled and dialysed against protein buffer (20 mM Tris pH7.4, 150 mM, NaCl, 10 mM glutamate). The purity was determined to >98% by SDS-PAGE analysis.

### Peptide spot array

Peptides covering the extracellular parts of γ2 and γ8 were spotted onto amino modified Whatman cellulose membranes (Figure 1B and Figure 1 – Figure supplement 1C) using a fully automatic Spot synthesizer (Intavis, Köln, Germany). The spot array consisted of hexameric overlapping peptides shifted by one residue. Peptide spotted membranes were rinsed with ethanol for 5 minutes, following three times 10 min washing with TBS and incubation with blocking buffer (Casein Blocking buffer (Sigma B6429), 150 mM Saccharose, in TBS) for 3 hours at RT. The blocking buffer was removed by three wash steps with TBS before the membranes were incubated overnight at 4°C with either 50 μg/ml protein (GluA2_LBD or GluA2_LBD_LY) in blocking buffer or blocking buffer only for control. Membranes were washed three times in TBS and incubated for 1.5 hours at RT with anti-poly_His Antibody (Sigma H1029) diluted 1:6000 in blocking solution followed by three washes (a’ 10 min) with TBS. Finally, membranes were incubated for 1.5 hours at RT with HRP-conjugated anti-mouse IgG Antibody (Sigma A5906; 1:1000 dilution in blocking buffer) and washed with TBS (three times a’ 10 min). Visualization of protein-binding was carried out using a chemo-luminescence substrate (Pierce™ ECL, ThermoFisher Scientific) and a Lumi-Imager™ instrument (Boehringer Mannheim, Germany). Spot-signal intensities were measured in Boehringer Light Units (BLU) and the software GeneSpotter 2.6.0 (MicroDiscovery, Berlin, Germany) was applied for data processing. Hits from peptides located within β-sheets were taken to be false positives, because when isolated these peptides likely form unphysiological β-sheets in a nonspecific manner with existing structures in the GluA2 LBD. To have an idea about reproducibility of this assay, we performed it twice with comparable results (source data is provided). The negative control showed no signal, indicating no unspecific binding of the anti-poly His to the peptides.

### Structural modeling

Initial γ2 and γ8 models were generated based on the crystal structure of claudin15 (PDB code: 4p79) using the SWISS-MODEL (42) and ProtMod server (part of the FFAS server, (43). Both models were incomplete (either lacking linker structures or failing to correctly trace transmembrane helix 3, TM3). Thus, we used COOT (version 0.8.7) to superpose the two generated models and to build the final model with an intact helix 3 and plausible extracellular loops 1 and 2. Superposing our final TARP models onto the γ2 molecules present in the AMPA-TARP cryo-EM structure (PDB code: 5kk2) in PyMOL (v1.6.0.0) yielded in the AMPA-TARP complexes shown in our Figures. The different possible orientations of Loop 1 were modeled using COOT. Unfortunately the LBD to TMD connecting linkers (S1-TM1 and S2-TM4) are not resolved in the AMPA-TARP cryo-EM structure. To better understand the Loop 2 participation in AMPA receptor regulation we used the crystal structure of GluA2 (PDB code: 3kg2) with resolved linkers and superposed it onto the receptor of our AMPA-TARP complex model (Figure 1 – Figure supplement 2). As the side chains of the possible interacting residues (507-QKS-510, 781KSK-783) located in the LBD-TMD linkers were not resolved in 3kg2 we modeled the most likely side chain conformations of these residues (Figures 7A and B). All figures were prepared with PyMOL or IGOR Pro.

## Acknowledgements

This work was funded by the Deutsche Forschungsgemeinschaft (DFG) – FOR 2518 (“DynIon”, to A.J.R.P.), the DFG Cluster of Excellence “NeuroCure” (DFG EXC-257, to ALC), and an Erwin-Schrödinger Postdoctoral Fellowship (J3682-B21) of the Austrian Science fund (FWF, to C.E). We thank Marcus Wietstruk and Ronny Schäfer for technical assistance.

## List of supplementary figures

- Figure 1 - Figure supplement 1. Loop interactions between TARPs and GluA2.
- Figure 1 - Figure supplement 2. The acidic patch on β4-TM2 of γ2 negatively modulates AMPA receptor gating.
- Figure 1 - Figure supplement 3. Sequence alignment of γ2 and γ8 constructs.
- Figure 2 - Figure supplement 1. Relief of polyamine block is not affected by loop mutations in γ2 and γ8.
- Figure 7 - Figure supplement 1. GluA2 linker mutants do not affect receptor kinetics or assembly with TARPs.

## List of source data files

- Figure 1 – Source Data 1: spot array quantitation
- Figure 1-Figure Supplement 2-Source Data 1: rectification indices for negative patch chimera
- Table 1 – Source Data 1: kinetics and steady state currents from electrophysiological recordings
- Figure 2 –Figure Supplement 1-Source Data 1: rectification indices for electrophysiological recordings of TARP chimeras
- Figure 6-Source Data 1: rectification indices for electrophysiological recordings of TARP deletion chimeras.
- Figure 7-Figure Supplement 1-Source Data 1: rectification indices for electrophysiological recordings of TARPs with GluA2 mutants.

## Supplementary Figures

**Figure 1- Figure supplement 1.**
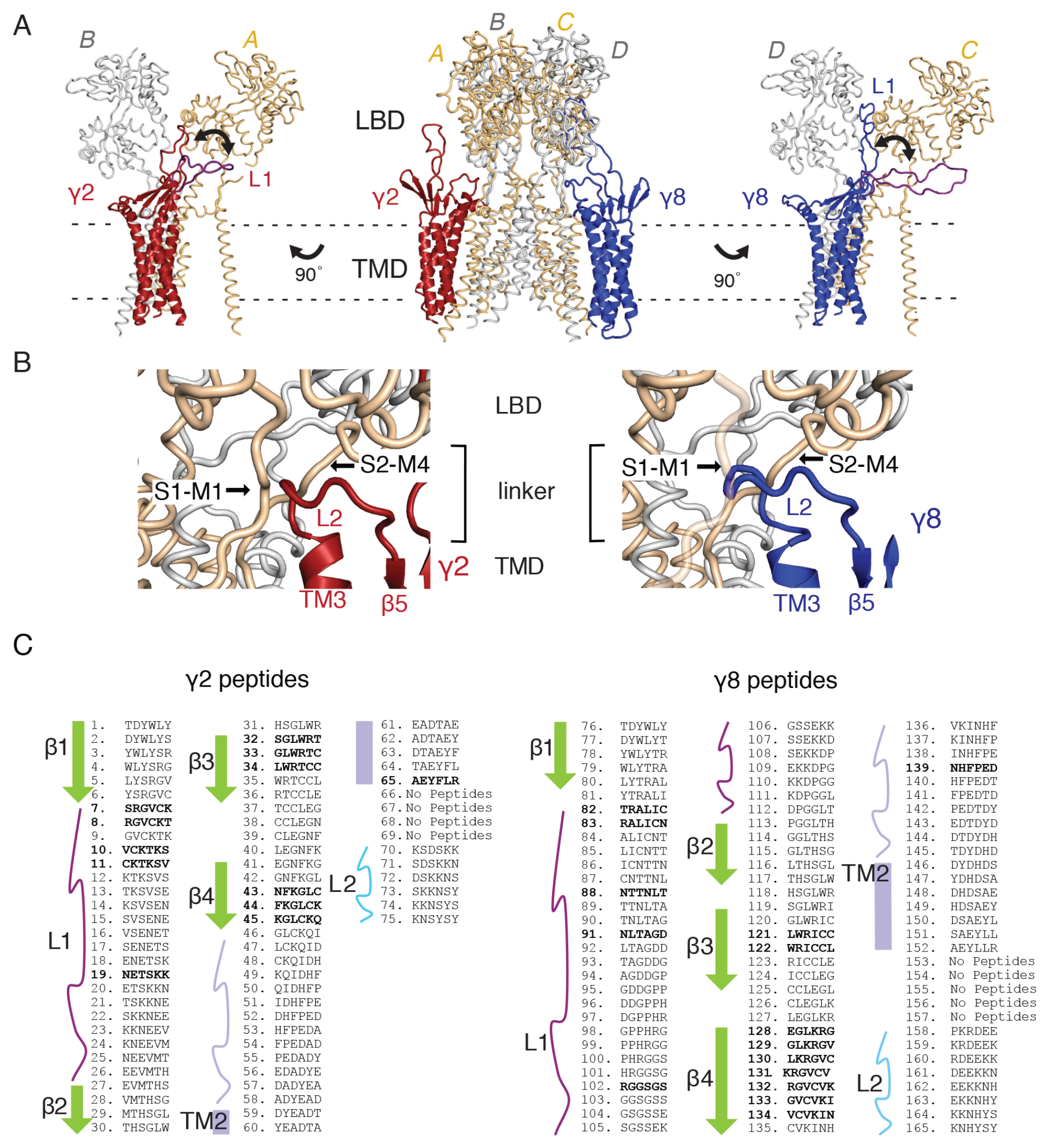
Loop interactions between TARPs and GluA2. **A)** The middle panel shows TARPs γ2 *(red)* and γ8 *(blue)* positioned between equivalent receptor subunits (A & B and C & D). We modeled L1 in two positions, either between the LBD dimer (colored as the respective TARP) or underneath the lower lobe of the LBD *(purple; left panel γ2, right panel* γ8). **B**) The model suggests L2 of both γ2 *(red, left panel)* and γ8 *(blue, right panel)* engages in similar interactions, independent from the TARP’s location in the complex. L2 is sandwiched between the receptor linkers (S1-M1 and S2-M4) connecting the LBD to the pore-forming TMD. C) Sequences of the hexameric TARP peptides used in the peptide mapping array are listed according to their position in the array. Secondary structure elements are shown in the same color code as in Figures 1B and C. Peptide sequences and quantitation are found in Figure 1–Source Data 1. Positive peptide hits are indicated as bold sequences.

**Figure 1- Figure supplement 2.**
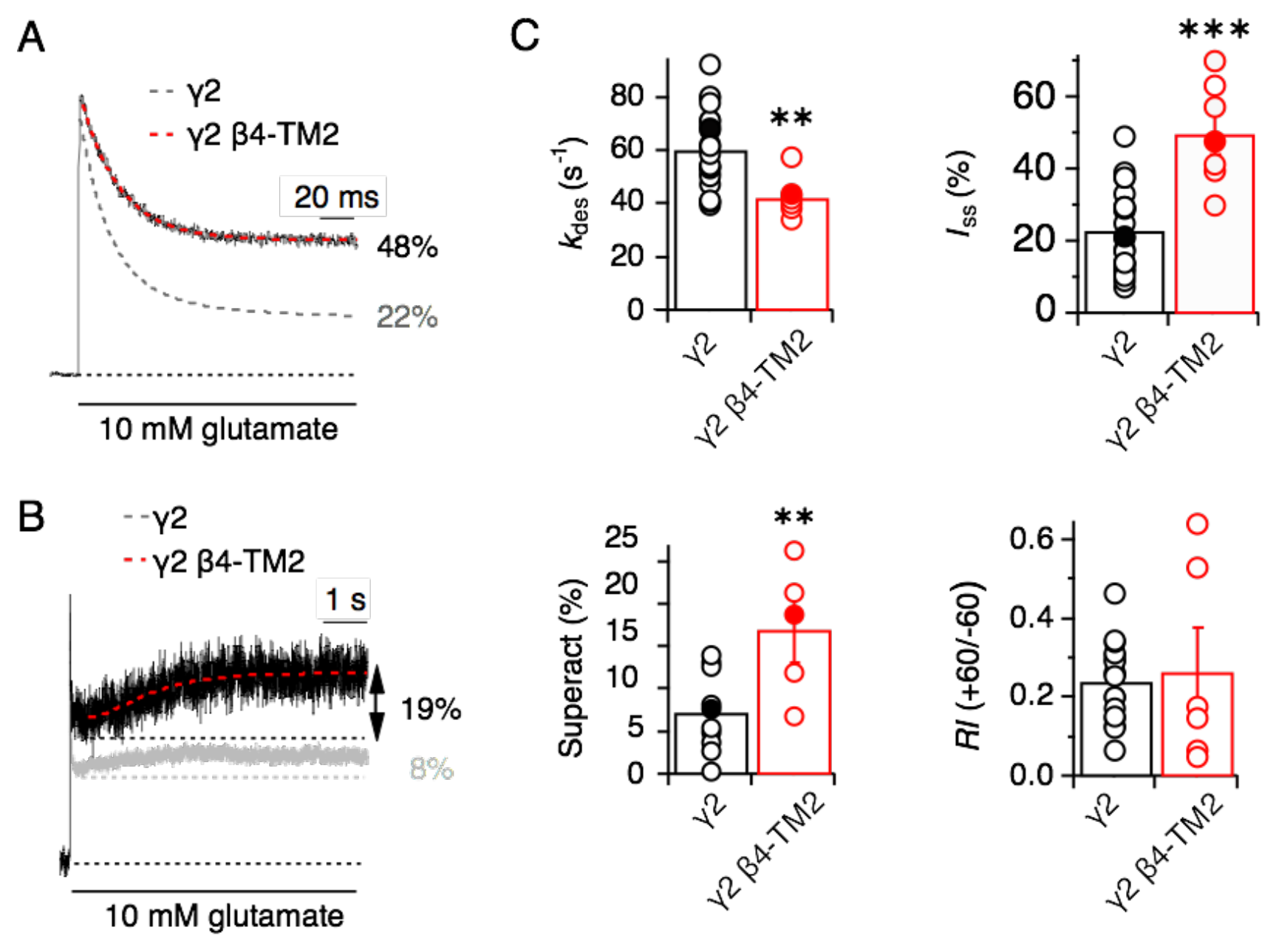
The acidic patch on β4-TM2 of γ2 negatively modulates AMPA receptor gating. **A**) Representative traces from γ2 β4-TM2 coexpressed with GluA2 *(red;* 3 negative charges removed) in response to a 500 ms pulse of 10 mM Glutamate show a substantial reduction in desensitization rate and extent (*k*_des_ = 43 s^−1^; *I*_ss_ = 48 %) compared to wild-type γ2 (dashed grey line). The mutations in γ2 β4-TM2 were D88G, E90S and D92G. **B**) Neutralizing the negative patch on the β4-TM2 increased γ2-mediated superactivation more than two-fold. The grey trace represents wild type γ2. **C**) Bar graphs showing the effects of neutralization of the γ2 negative patch on desensitization, steady-state current, superactivation. The rectification index was not changed, indicating relief of polyamine block was intact. Filled symbols correspond to the traces shown in A) and B). \*\**p* < 0.01, \*\*\**p* < 0.001, against γ2. Source data for kinetic data in panel C is found in Table 1 – Source Data 1. Source data for rectification indices in panel C is found in Figure 1 – Figure Supplement 2 – Source Data 1. Error bars represent s.e.m.

**Figure 1- Figure supplement 3.**
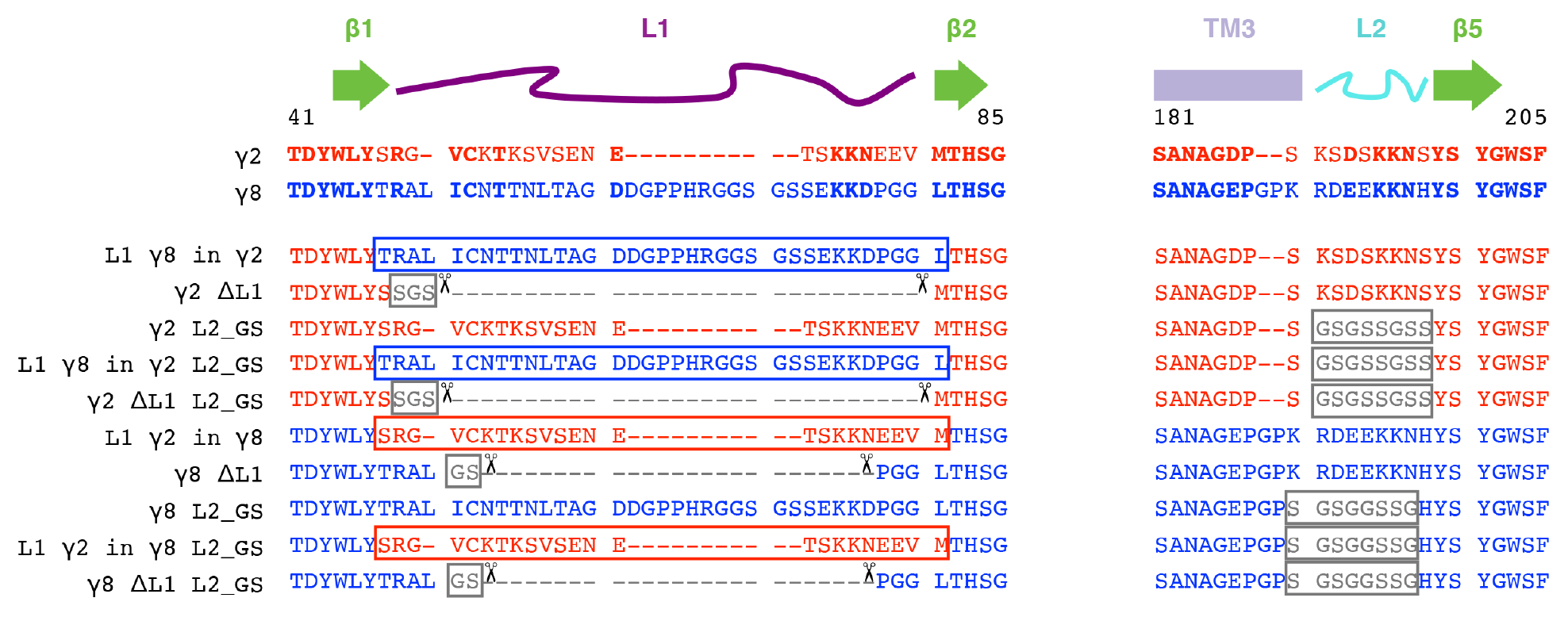
Sequence alignment of γ2 and γ8 constructs. The sequences of the extracellular regions Loop1 (L1, *purple)* and Loop2 (L2, *cyan)* of γ2 *(red)* and γ8 *(blue)* are aligned with the secondary structural elements on top. Constructs carrying deletions (L1, indicated by scissors), neutralization (L2, glycine-serine (GS)-Linker) and chimeras (switching L1 between the TARPs) and combinations of these are shown below.

**Figure 2 Figure supplement 1.**
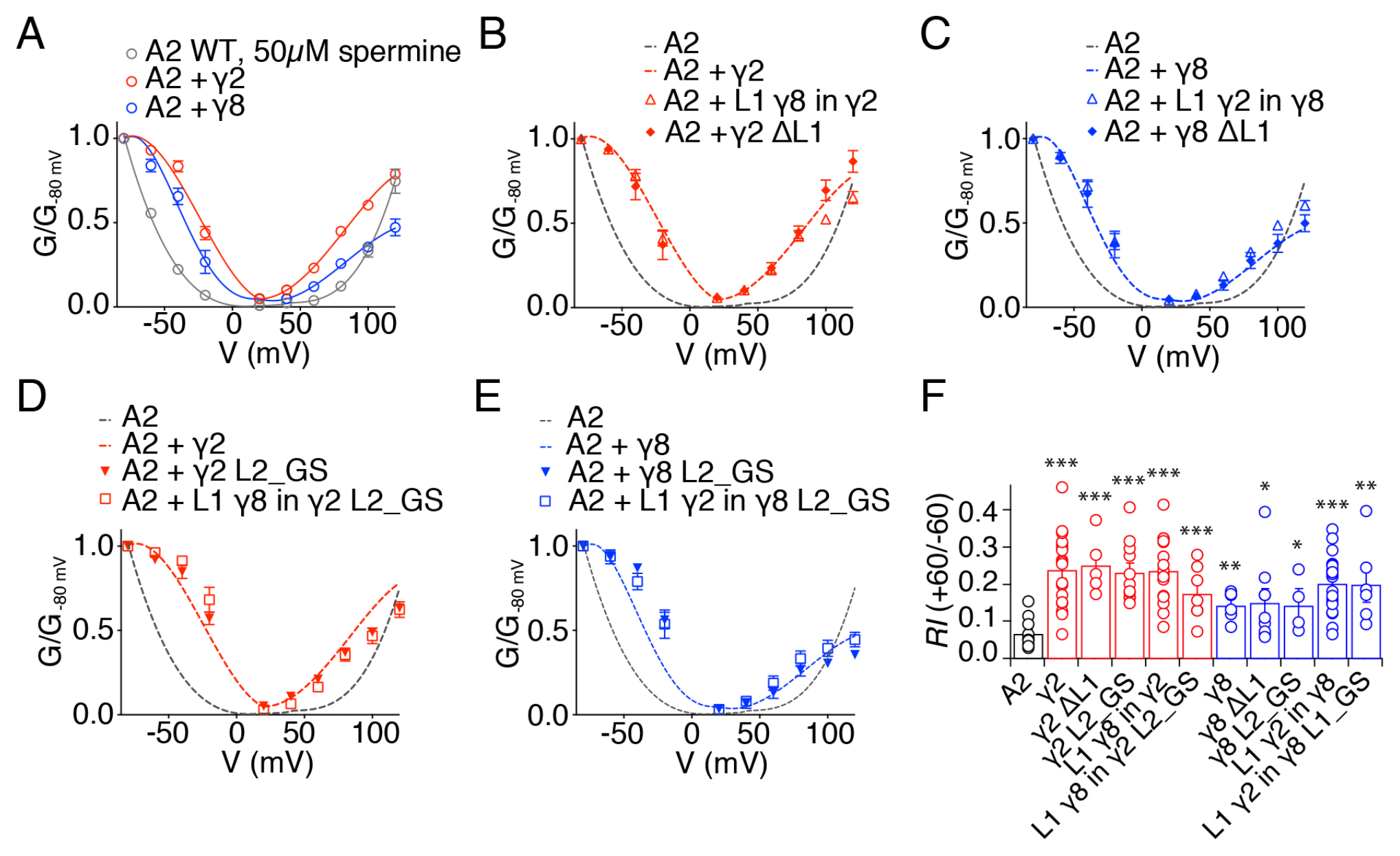
Relief of polyamine block is not affected by loop mutations in γ2 and γ8. **A**) Normalised conductance-voltage plots show that TARP γ2 (red) is better at relieving the polyamine (PA) block of unedited GluA2 receptors *(grey)* than γ8 (*blue*). **B**) Relief of PA block by γ2 L1 mutants *(filled symbols)* is indistinguishable from that of the wild type construct *(dashed line)*. GluA2 WT is shown in grey for comparison. **C**) Replacing L1 of γ8 with that of γ2 or deleting it does not affect its ability to relief PA block of GluA2(Q) receptors. **D**) Neutralizing L2 in γ2, alone or in combination with L1 from γ8, does not affect PA block. **E**) Neutralizing L2 in γ8 and in γ8 with L1 from γ2 show similar PA block relief as γ8 wild type. **F**) Bar graph summarizing the rectification index (*RI*, calculated as the ratio between the current recorded at +60 mV and that recorded at -60 mV) of γ2 (in *red)* and γ8 (in *blue)* loop mutants coexpressed with GluA2(Q). Currents were recorded in the presence of 50 μM spermine in the pipette solution. * *p* < 0.05, ** *p* < 0.01, *** *p* < 0.001, against GluA2(Q). Source data for panel F is found in Figure 2 – Figure supplement 1 – Source Data 1. Error bars represent s.e.m.

**Figure 7-Figure supplement 1.**
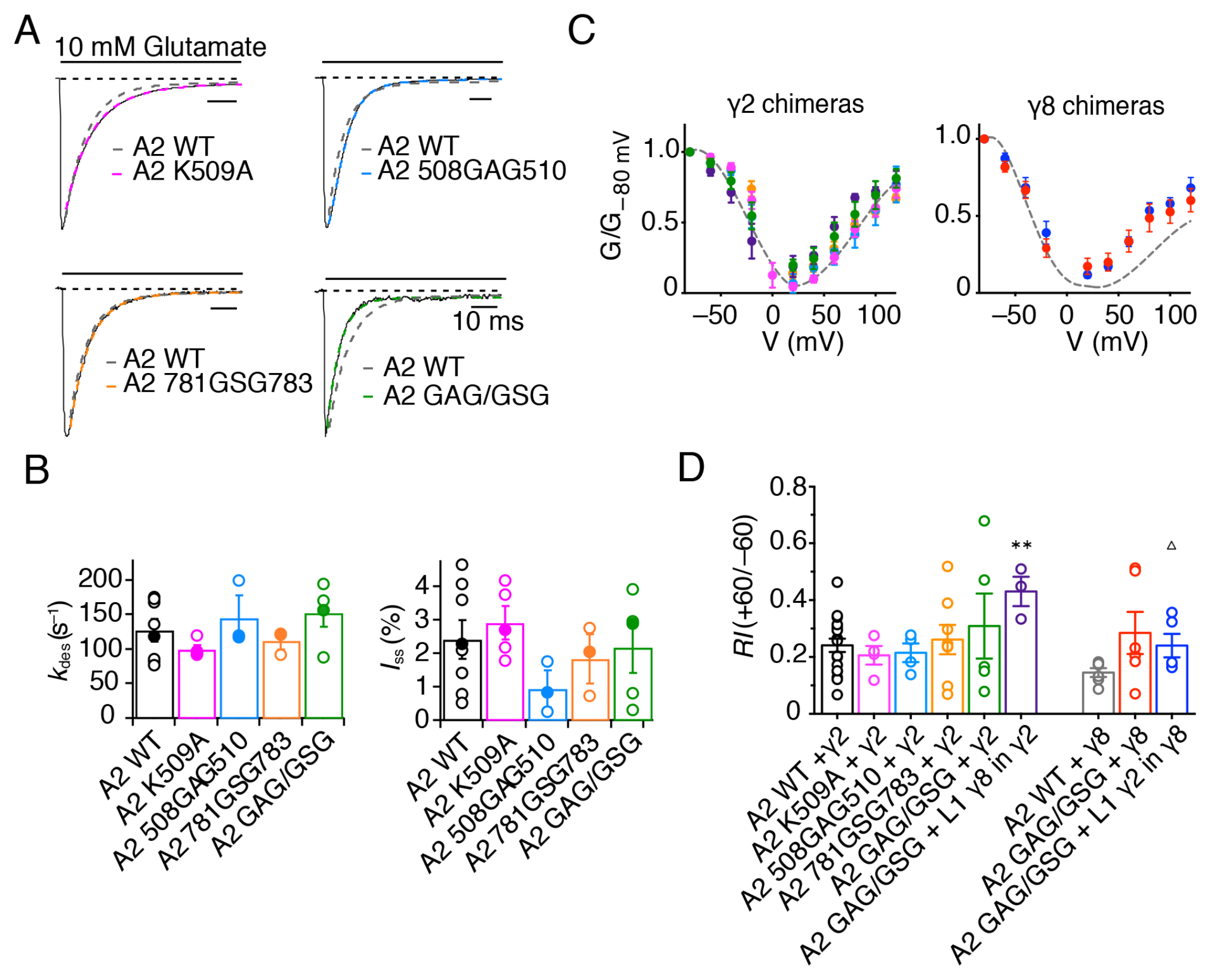
GluA2 linker mutants do not affect receptor kinetics or assembly with TARPs. **A**) Representative traces from GluA2 linker mutants in response to 500 ms pulses of 10 mM Glutamate. GluA2 WT is shown in grey. **B**) Bar graph summarizing the desensitization kinetics and the level of steady state current for GluA2 linker mutants. Filled dots represent the traces shown in A). **C**) **and D**) GV responses and rectification index for GluA2 mutants in complex with γ2 WT or L1 γ8 in γ2 chimera *(left)* and γ8 WT and L1 γ2 in γ8 chimera *(right)*. ** *p* < 0.01, against γ2; Δ *p* < 0.05, against γ8. Source data for panel C is found in Table 1 – Source Data 1 and source data for panel D is found in Figure 7 – Figure supplement 1 – Source Data 1. Error bars represent s.e.m.

